# 14-3-3β paralog is inhibited by acetylation during differentiation to the osteogenic lineage

**DOI:** 10.1101/2019.12.17.879155

**Authors:** Yesica R Frontini-Lopez, Aldana D Gojanovich, Marina Uhart, Diego M Bustos

**Affiliations:** Laboratorio de Integración de Señales Celulares, Instituto de Histología y Embriología de Mendoza (IHEM-CONICET-UNCuyo), 5500, Mendoza, Argentina; CReM, Boston University School of Medicine and Boston Medical Center, 02118, Boston, MA, USA; Facultad de Ciencias Exactas y Naturales. Universidad Nacional de Cuyo, 5500, Mendoza, Argentina

## Abstract

14-3-3 protein family binds and regulate hundred of serine/threonine phosphorylated proteins. Considered as redundant, ubiquitous and constantly expressed this protein family was treated as an accessory for many signaling systems. Here we studied the reversible inhibition by acetylation of its essential N-ε-lysine 49/51 residue during the osteogenic differentiation of human adipose-derived stem cells (ASC). We found that during the differentiation of ASC the levels of 14-3-3 acK49/51 increase showing that inhibition of 14-3-3 is necessary for this process. Among the 7 paralogs of this family, the inhibition by this posttranslational modification occurs mostly on the paralog β located specifically in the nucleus where 14-3-3 was described to binds to H3 histone and many transcription factors. Short hairpin RNA silencing of 14-3-3β gene but not 14-3-3γ increases significantly the osteogenic potential of the cells. These results show that specifically 14-3-3β is a negative regulator of osteogenesis and its inhibition by acetylation on lysine 51 is the cellular mechanism to regulate it.

## Introduction

Adipose-derived stromal/stem cells (ASCs) are naturally multipotent, being able to commit to various lineages such as adipogenic, osteogenic, and chondrogenic [1]. Osteogenesis is the process by which new bone is synthesized [2], however, the available data on osteoblast differentiation is fragmented and incomplete. A major source of discrepancies in this research area is the use of cell lines instead of primary cells [3]. Several cellular pathways, as Hippo, Wnt, TGF-β, BMP and Notch, are known to be involved in this differentiation process. They all converge to the Runt-related transcription factor 2 (RUNX2), the master transcriptional regulator of osteoblast differentiation [3]. RUNX2 is predominantly a fetal factor, while in the adult organism its levels are low. This transciption factor regulate the expression of important genes for mineralization, including osteopontin, bone sialoprotein, osteocalcin, osteoprotegerin, Rankl, and many others [3,4]. Recent studies show that other signaling pathways like JNK, HedgeHog, and ERK1/2 pathways are also involved in the process [1,5]. Epigenetic mechanisms mediated by histone acetylases (HAT) and deacetylases (HDACs) have been shown to be associated with osteoblast differentiation and maturation [6]. The first identifiable substrates for HDACs are the histone N-term Lys residues [7], and their activity is usually directed by the formation of protein complexes. For example, Westendorf *et al.* showed that HDAC6 (a class IIb deacetylase) binds to RUNX2 in differentiating osteoblasts and this interaction is necessary for the repression of the p21 promoter in preosteoblasts [8]. Another set of evidence of HDAC participation in the osteogenic differentiation came from the use of general and class specific inhibitors of HDAC. The use of HDAC inhibitors *in vivo* and *in vitro* models have shown that this protein family have multiple and intricate roles during osteoblast maturation, including inhibition of proliferation and induction of differentiation [9,10]. The HDAC inhibitor sodium butirate (NaBu) for instance, regulates the expression of osteoblast-specific genes enhancing RUNX2-dependent transcriptional activation in MC3T3-E1 preosteoblast cells [11]. However, the precise mechanism and all the components in this machinery have not been fully elucidated yet. Recently, it has been shown that modification of Lys residues by acetylation is in some way related to the cellular mechanisms of phosphoryaltion [12]. The HDACs 4, 5, 7, and 9 (all these belonging to class IIa) contain N-terminal extensions with binding sites for the phospho-readers 14-3-3 proteins. The 14-3-3 protein family sense and transduce signaling information by binding to serine or threonine phosphorylated proteins [13]. This protein family mediates the regulation of many signaling pathways, including those related to osteogenesis as Hippo, TGF-β, IGF1R, PI3K/AKT, and Wnt/β-catenin among others [13]. The cross-talk feedback between the signaling through protein phosphorylation and acetylation is given by the fact that 14-3-3 proteins can be inhibited through acetylation on the essential Lys49/51 (numbers depend on the paralog) by -at the moment-an unknown acetylase [14] and deacetylated by a class IIb HDAC, the HDAC6 [15]. This process is known as “regulation of regulators”, as phosphoreaders domains-which can bind phosphosites and regulate the bearing protein-are in turn regulated by acetylation [16]. The 14-3-3 protein family is composed by 7 paralogs in mammals; among which specific members have been reported as either positive or negative regulators of osteogenesis. The downregulation of 14-3-3β in calvaria organ cultures resulted in increased bone formation [17], whereas 14-3-3ε in exosomes released by osteoblasts and osteocytes in response to dynamic compression has positive paracrine effects [18]. Interestingly, TAZ, a known mechanosensor and transcriptional modulator of the Hippo pathway, has also been linked to 14-3-3 proteins, which regulate its nuclear-cytoplasmic shuttling [19], indicating an additional role for 14-3-3 proteins in bone signaling mediation.

Here, we studied the Lys49/51 actylated state in 14-3-3γ and β paralogs in ASC isolated from human donors. We found that acetylation of Lys49/51 by the MYST family acetylase HBO1 has dramatic effects on 14-3-3 sub-cellular activity localization in general, and in the activity regulation of β and γ paralogs in particular during the process of osteoblast differentiation.

## Results

### 14-3-3 paralogs are regulated differently during ASC osteogenic differentiation

Previous results from our laboratory showed that during the differentiation of NIH3T3-L1 preadipocytes cell line, the expression levels of 14-3-3 protein family paralogs are independently regulated [20]. As it has been proposed that adipogenic and osteoblastic differentiation are highly related phenomena, we analyzed the levels of six 14-3-3 paralogs in ASC treated with specific osteogenic differentiation medium (ODM, for composition, see Material and Methods section). The ACS were obtained from healthy patients who underwent voluntarily a dermolipectomy surgery, and donated abdominal subcutaneous adipose tissue after signing bioethics committee-aprooved informed consent documents [21]. After isolation, ASC were characterized by flow cytometer analysis [21], using a panel of three positive (CD105, CD90 and CD73) and five negative (CD45, CD34, CD11b, CD19 and HLA-DR) markers as recommended by [22] and differentiation to mesodermal linages (Fig S1). Figure 1 shows the levels of 6 of the 7 paralogs of 14-3-3 proteins expressed in ASC, the paralog SFN/σ has a more restricted expression pattern [23] and was undetectable in our samples. Among the other 6 paralogs, 3 of them showed differences in expression during the process of osteoblast differentiation (Fig 1A). We observed that the β and ε paralogs increased their levels of expression whereas the γ paralog decreased its levels after 21 days of osteogenic differentiation. In previous results from our laboratory, we observed an opposed tendency in expression of the 14-3-3 γ paralog during the differentiation of NIH3T3-L1 cells to adipocytes [20] and for both γ and β paralogs in ASC (manuscript in preparation). About the 14-3-3 ε paralog, its role in bone and cartilage formation was already proposed, as a soluble mediator in the communication between subchondral bone and cartilage [18,24]. This paralog was also detected as a component in several exosomes, and it secretion and extracellular activity was described [25]. We believe that its role in osteogenesis could be of a completely different nature than β and γ paralogs. For these reasons, we focused on elucidating β and γ 14-3-3 paralogs roles in osteogenic differentiation of ASCs.

**FIGURE 1.**
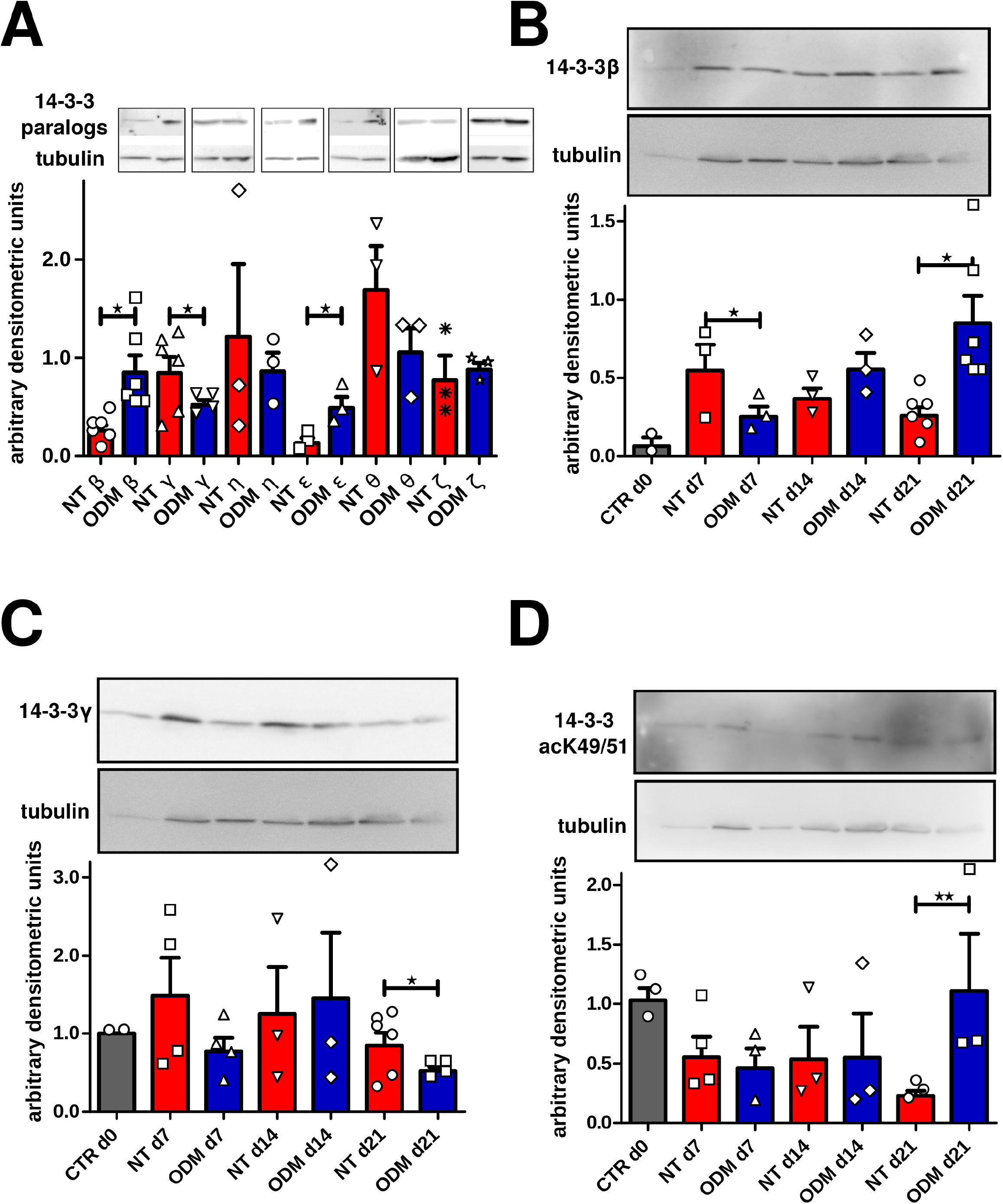
Western blot analysis of 14-3-3 proteins relative levels in non-treated and ODM-treated ASC. A) Total cell lysates were prepared at day 21 after osteogenic differentiation induction, 40 μg were loaded per well and separated by SDS-PAGE. Blots were divided for reaction with specific antibodies against each of the different 14-3-3 paralogs and α-tubulin as loading control. Bars in the graphs represent mean ±SD, of values from 3 experiments quantified by densitometry and expressed relative to that of tubulin values from the same Western blot line. Asterisks indicate significant differences between samples (using the Wilcoxon rank-sum test, *p* < 0.05). B) Same as in A, except that lysates were prepared at 7, 14 and 21 days (d7, d14 and d21 respectively) after osteogenic differentiation induction and that blots were incubated with antibodies against 14-3-3β and α-tubulin. C) Same as in B but using antibodies against 14-3-3γ. D) Same as in B but using antibodies against 14-3-3 acK49/51. A to D) Data were similar in two other experiments.

To further characterize the pattern of expression of β and γ paralogs in ASC differentiating process to osteoblasts, we performed a 21 days time course expression. We observed, in the case of β paralog, a strong induction on its expression at 7 days of growing in supplemented DMEM with a slight decrease at day 21 (Fig 1B). However, when ASC were induced to differentiate to osteoblasts by the addition of ODM drug cocktail, we observed an opposed expression pattern of the 14-3-3β paralog (Fig 1B). The addition of ODM medium to ASC reduces the levels of 14-3-3β at the beginning of the differentiation, a pattern that is reverted during the course of the differentiation (Fig 1B). In the case of 14-3-3γ paralog, its expression during the osteogenic differentiation does not follow a clear tendency, but its level after 21 days of differentiation were the lowest observed (Fig 1C).

### Inactivation of 14-3-3 proteins by acetylation of Lys49/51 increases during ASC osteogenic differentiation

HDAC inhibitors, including NaBu, have a role regulating the expression of osteoblast-specific genes like RUNX2 [11], and others [1]. The acetylation of Lys49/51 in 14-3-3 proteins produce an inactive protein, avoiding the binding to their targets [13,14]. The Lys49/51 is essential for the binding because it forms one of the three hydrogen bonds in the final complex between 14-3-3 and the phosphorylated Ser/Thr in 14-3-3’s partners (Fig S2A and B) [13]. In this sense it appeared interesting to measure not only the levels of individual 14-3-3 paralogs but to also measure the levels of acetylated Lys49/51 in these proteins. To do this, we used a specific 14-3-3 acK49/51 antibody, which was extensively characterized in our laboratory (Fig S3). Figure 1D showed the levels of 14-3-3 acK49/51 during ASC osteogenic differentiation. We observed that the levels of inactivated 14-3-3 slightly decreased during the course of the experiment when the cells were growing in complete DMEM. However, when the cells were treated with ODM drug cocktail, after 21 days of differentiation the levels of inactive 14-3-3 were significantly higher than the levels on nontreated ASC. This suggests that acetylation of 14-3-3 proteins could have a biological role during ASC osteogenic differentiation.

### Subcellular localization of acK49/51 14-3-3 proteins, and HBO1 acetyltransferase

The 14-3-3 proteins are highly and broadly expressed in the cell [23]. We studied the subcellular localization of 14-3-3 acK49/51 during the ASC osteogenic differentiation process to observe possible enrichments in specific subcellular compartments. It has been theoretically proposed that different subcellular pools of 14-3-3 could be affected by site-specific acetylation [26]. Figure 2 shows immunefluorescence (IFI) experiments of ASC at day 21 of osteogenic differentiation. Nuclear staining of 14-3-3 acK49/51 was clearly observed in both conditions, non-treated (NT) and ODM treated cells. However the later showed significantly higher signal, and it is also possible to observe cytoplasmatic staining. Lysine acetyltransferases are mostly located in the nucleus, where their main substrates are histones. However many HATs, such as members of the CBP/p300 and MYST families, have been shown to acetylate nonhistone proteins [27]. Among the MYST family (KAT5, 6A and B, 7 and 8), only the KAT7 (HBO1) is reported to be detected in the cellular cytoplasm, information recovered from Uniprot database [28]. As shown in figure S2C, the catalytic site is highly conserved in the MYST family, and target specificity has not been reported yet. We decided to test HBO1 for 14-3-3 protein acetylation. In figure 3A we show the results of 14-3-3acK49/51 and HBO1 acetyltransferase IFI experiments of ASC during their differentiation to osteoblasts from day 0 to day 21. Images were quantified by using a python scripts to show the enrichment in nuclear localization for 14-3-3acK49/51 for many conditions (Fig 3B). This nuclear staining for 14-3-3acK49/51 is present at all days and conditions (NT and ODM), however in the later we also observed nuclear speckles, which resemble liquid-like condensates [29]. The cytoplasmic staining was more evident at day 14 and remains until the end of the treatment on day 21. Although we did not expected to find high degrees of colocalization because our hypothesis was that HBO1 and 14-3-3 could be an enzyme-substrate pair, we analyzed the images using 3 different colocalization analysis methods (Fig S4). Depending on the stage of the differentiation process, we found complete colocalization with different intensities, or partial colocalization. In most of the cases the superposition of molecule A (14-3-3 acK49/51) to molecule B (HBO1) was greater than the superposition of B to A. This could be an effect of the endogenous expression levels of each molecule or that HBO1 was acetylating other proteins.

**FIGURE 2.**
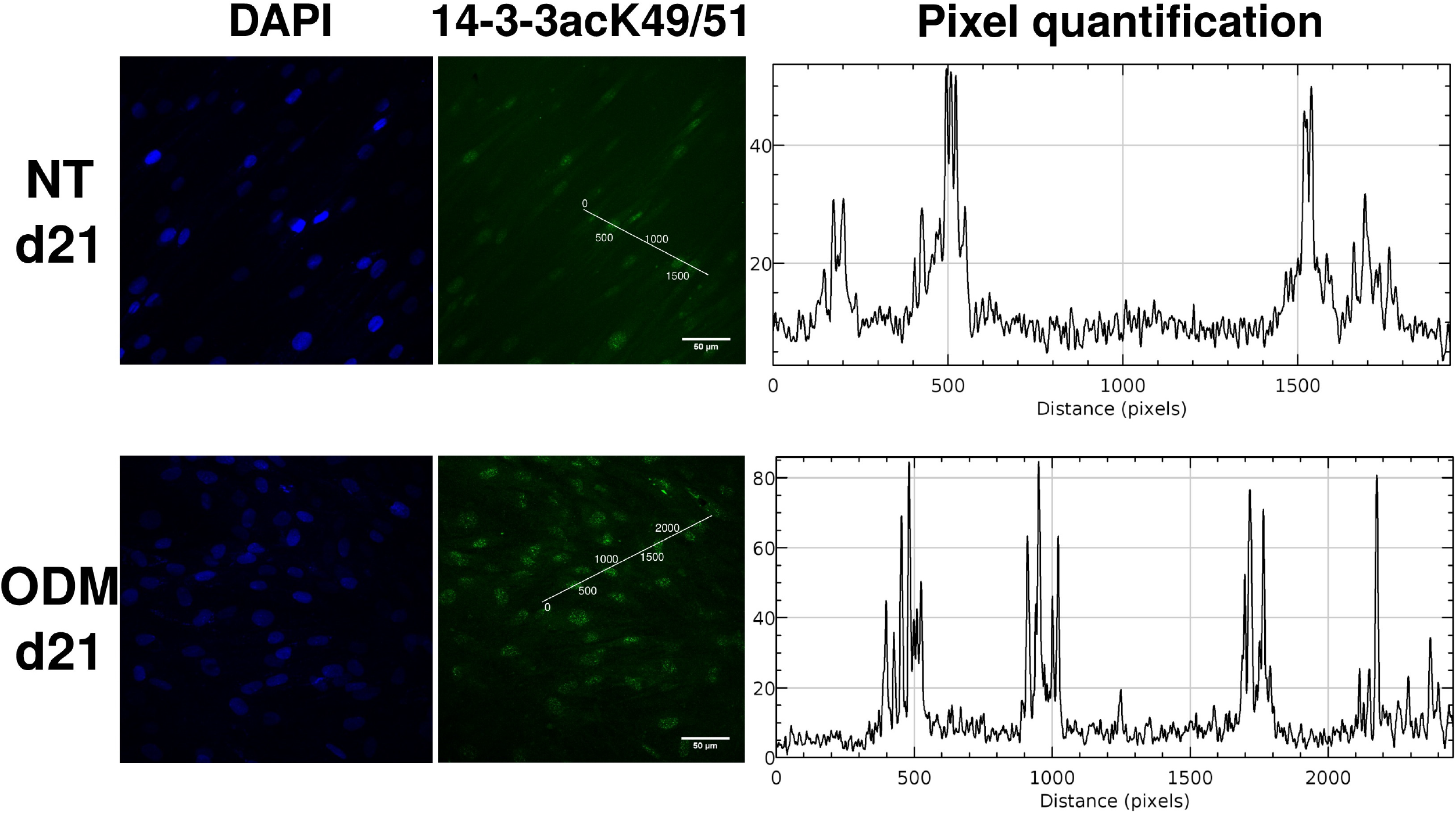
14-3-3 acK49/51 localization in ASC induced to the osteogenic lineage. Confocal images of 14-3-3 acK49/51 in non-treated (NT) and ODM treated ASC (magnification 40X). Cells were fixed at day 21 after growing in complete DMEM (NT) or osteogenic differentiation medium (ODM), and an Indirect Immunofluorescence (IFI) was performed as in [21]. Green signal corresponds to 14-3-3 acK49/51 proteins detected by IFI (Alexa Fluor 488); blue, DAPI; images were quantified to show localization of the immunoreactivity. Data were similar in two other experiments.

Figure 4 shows an increase of the 14-3-3acK49/51 levels in response to the treatment with the inhibitor of lysine deacetylases NaBu (A) or to the transfection with the HBO1 human gene (B). This results suggested that HBO1 acetyltransferase acetylates 14-3-3 on its Lys49 residue (or 51 depending on the paralog), although other acetyltransferase/s could also carry out this PTM due to the low specificity of this enzyme family.

It has been proposed that NaBu regulates osteogenesis in human amniotic mesenchymal stem cells and other stem cells [30]. We treated ASC with NaBu at a final concentration of 0.5 mM for three days before the induction of the osteogenic differentiation. After the treatment for 3 days with NaBu, the cells were incubated with complete DMEM (NT) or ODM to induce the differentiation to osteoblasts. It was observed a significant increment in the cytoplasmic staining of 14-3-3acK49/51 for all conditions and times (Fig 4). This result suggests that inhibition of 14-3-3s by acetylation of their critical Lys49/51 is part of the osteogenic differentiation program in ASC.

Immunoreactive HBO1 and 14-3-3acK49/51 were detected in the cytosol and the nucleus (Fig 3 and 4 and S5). To identify if the cytoplasmic localization of both HBO1 and 14-3-3acK49/51 is related to their final degradation, the cells were treated with the proteasome inhibitor lactacystin (lactacys 10 μM) [31]. The results show that in our conditions neither HBO1 nor 14-3-3acK49/51 have signals of cytosolic degradation (Fig. S5).

**FIGURE 3.**
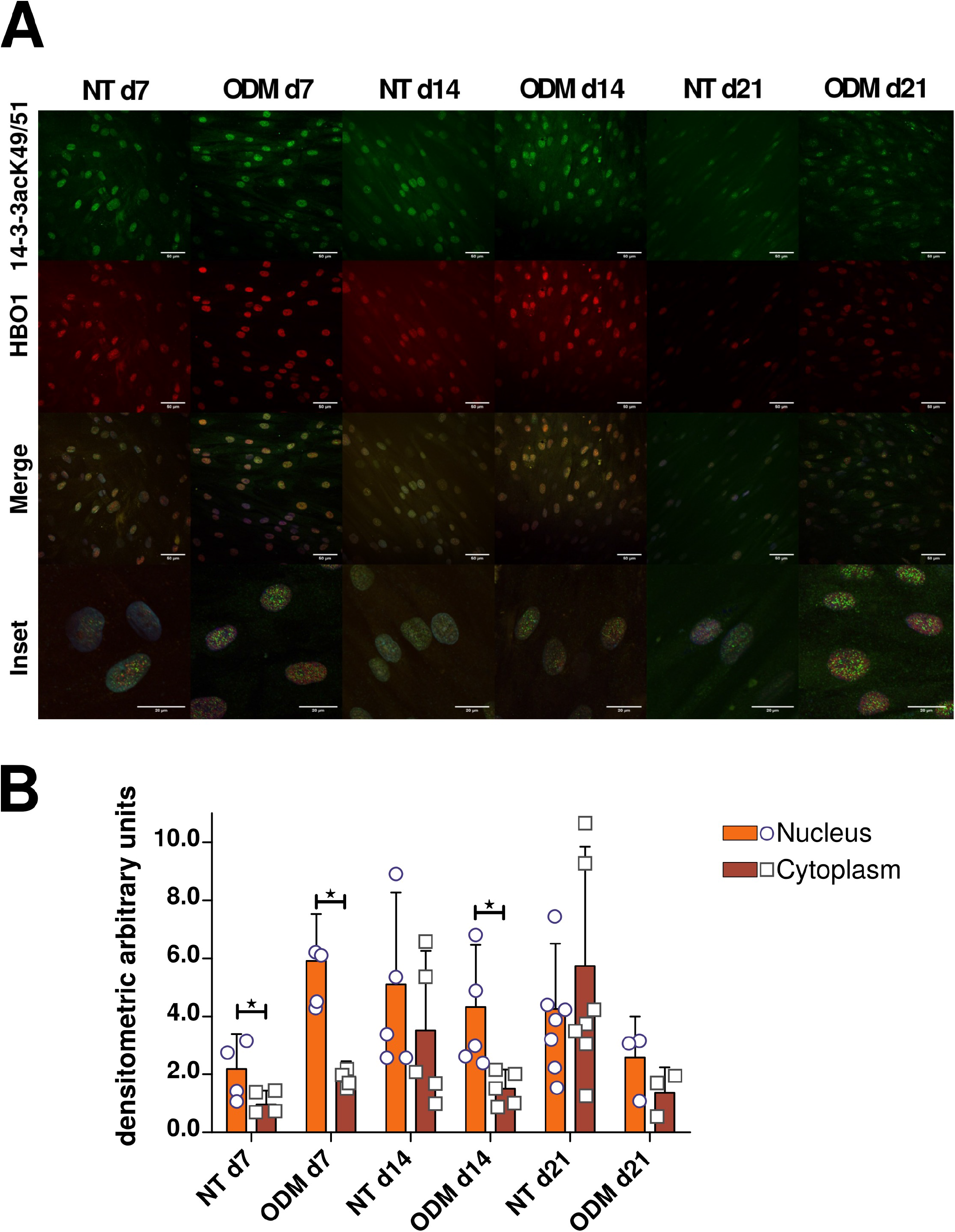
14-3-3 acK49/51 and HBO1 localization in ASC induced to the osteogenic lineage. Same as Fig. 2 except that ASC were fixed at different days (d7, d14 and d21) after growing in complete DMEM or osteogenic induction. Green and blue, same as in Fig. 2; red signal corresponds to HBO1 detected by IFI (Alexa Fluor 594). Data were similar in two other experiments.

**FIGURE 4.**
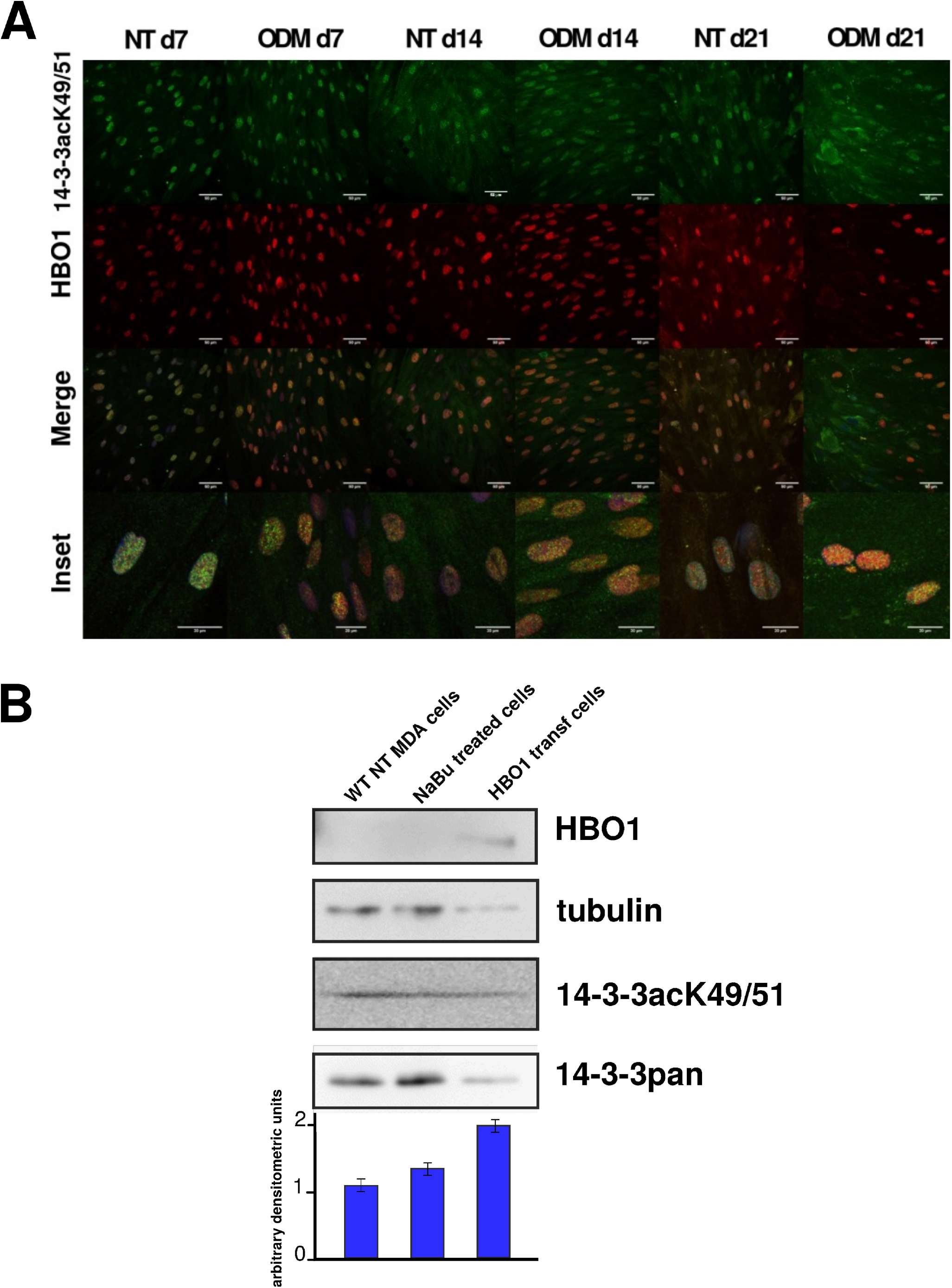
14-3-3 acK49/51 and HBO1 in NaBu treated cells. A) 14-3-3acK49/51 protein levels analyzed by Western blot after three days treatment of MDA-MB231 cells with the inhibitor of lysine deacetylases NaBu (0.5 mM) or 48 h after the transfection with the HBO1 human gene. B) Same as Fig. 3 except that ASC were treated with 0.5 mM NaBu for three days before induction of the osteogenic differentiation. Magnification 40X. A and B, data were similar in two other experiments.

### 14-3-3 paralog β-but not γ-is regulated by acetylation in Lys49/51 during ASC osteogenesis

We were interested in determining the levels of inhibition by acetylation of the specific 14-3-3 paralogs β and γ. To do this, we performed immunoprecipitation (IP) experiments by using specific and validated 14-3-3 paralog antibodies in sera [32]. These antibodies were developed in the laboratory of Dr Aitken who kindly donated to us. These antibodies have been extensively validated in many laboratories, including our [20,21,33]. In figure 5 we show the IP experiment result in which we used the anti-14-3-3β (and the corresponding control, pre-bleed serum) to immunoprecipitate NT and ODM ASC lysates followed by the detection using a commercial 14-3-3 pan antibody and the specific 14-3-3acK49/51. The results showed that 14-3-3β is specificly acetylated at its Lys51 when ASC were induced to differentiate to the osteoblast lineage (Fig 5A). No signal was detected when the same amount of pre-bleed serum was used in the IP. When we used antibodies against 14-3-3γ we observed that the levels of this paralog and its Lys49 acetylated form were lower in ODM treated ASC compared to the levels in NT ASC at d21 of the experiment (Fig 5C). These results showed that acetylation of Lys51 on β and Lys50 on γ paralog are regulated during the differentiation of human ASC to osteoblasts.

**FIGURE 5.**
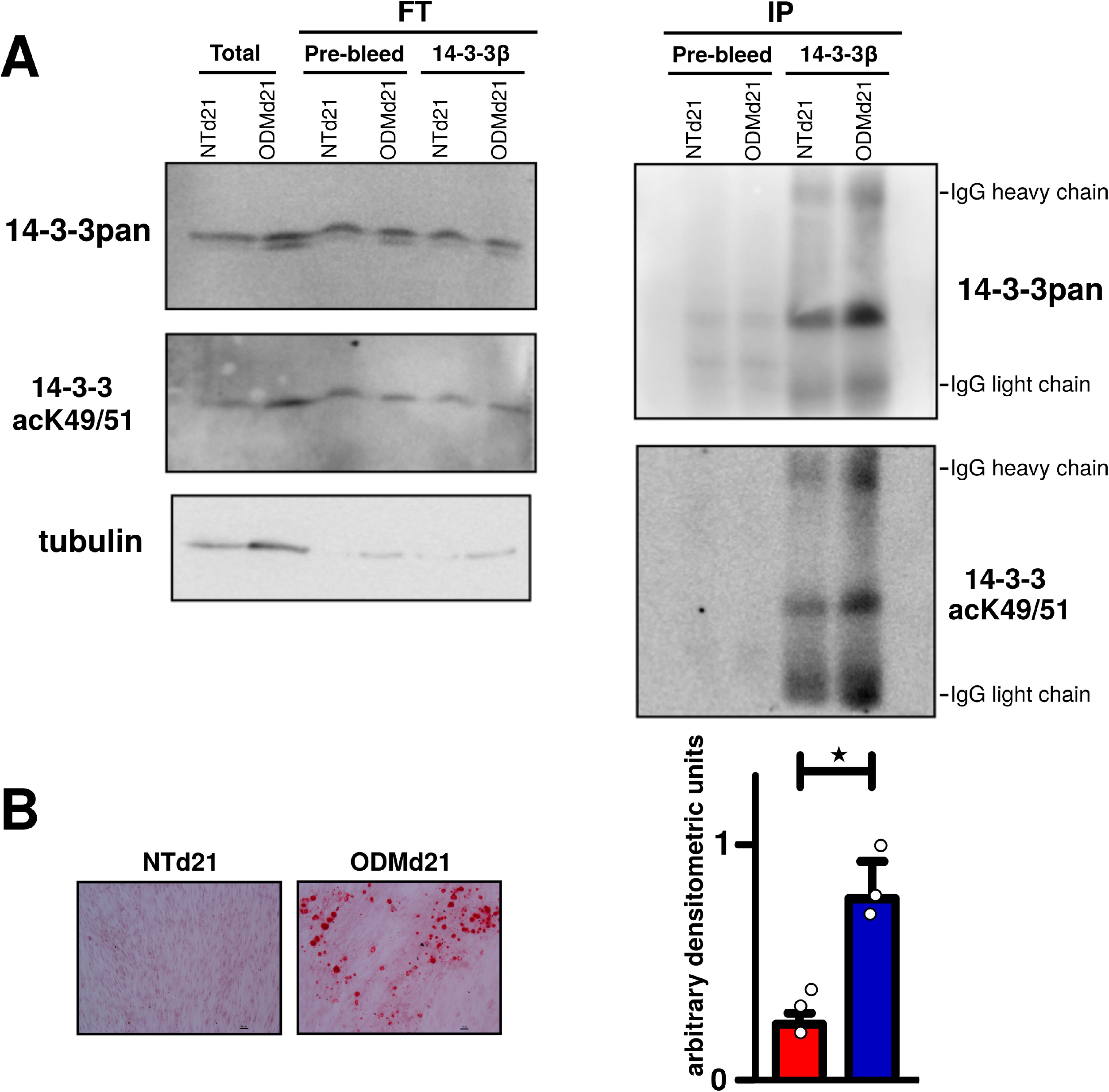
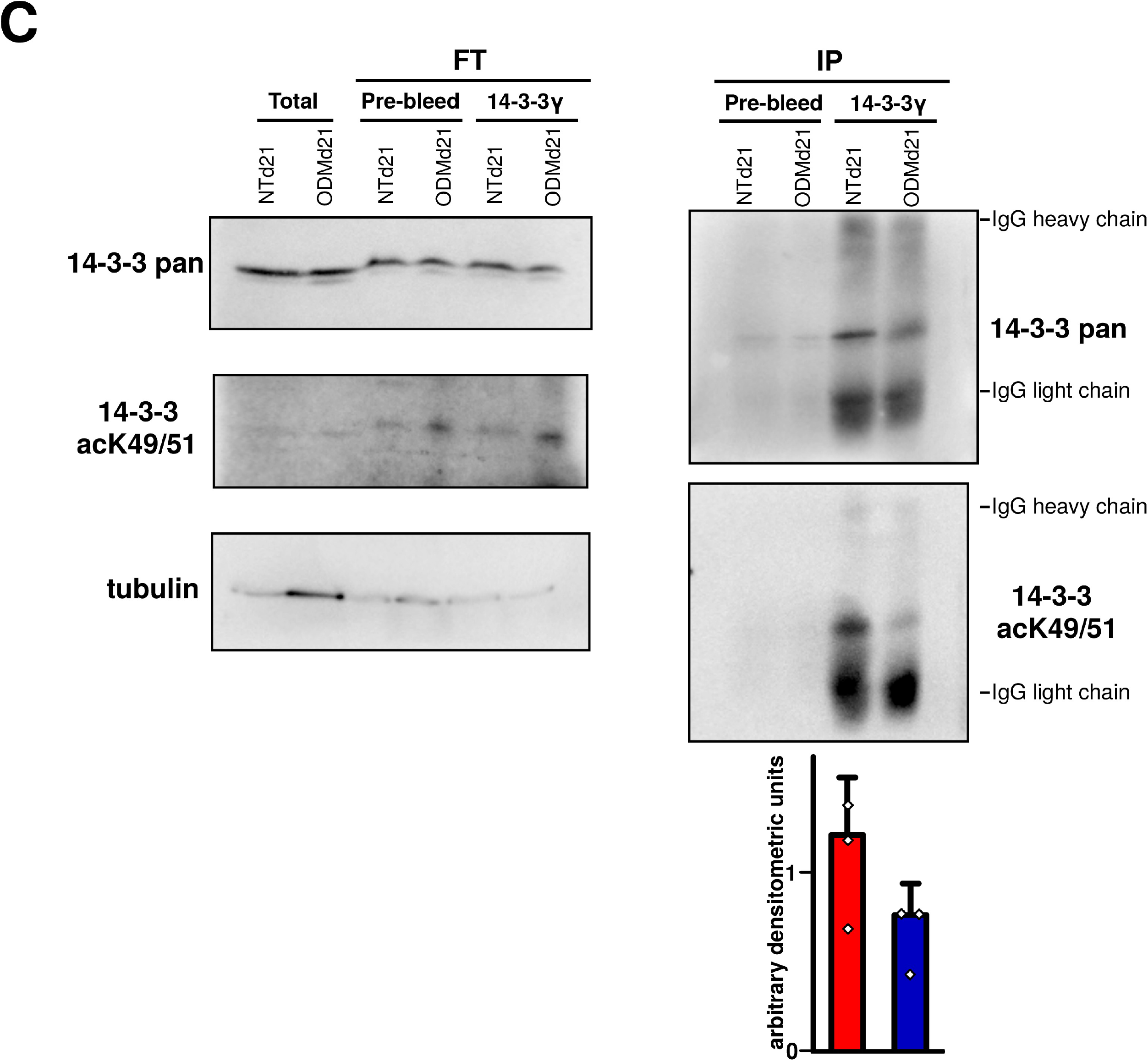
Immunoprecipitation of 14-3-3β and γ paralogs from ASC induced to the osteogenic lineage. After 21 days of osteogenic differentition, total cell lysates samples (250 μg of total protein) were immunoprecipitated with pre-bleed serum (nonspecific antibodies) or specific antibodies against 14-3-3β (A) and γ (B) paralogs. Samples (20 μl) of total cell lysates (Total), flow through (FT) or immunoprecipitated proteins (IP) eluted from beads in 100 μl of gel-loading buffer were separated by SDS-PAGE, and blots were divided for reaction with antibodies against 14-3-3 pan, 14-3-3 acK49/51 and tubulin. Bars in the graphs represent mean ± SD, of values from 3 experiments quantified by densitometry and expressed relative to that of 14-3-3 pan values from the same Western blot line. Data were similar in two other experiments.

### shRNA 14-3-3β and 14-3-3γ change the transdifferentiation potential of NIH3T3-L1 cells in opposite ways

We examined the effect of shRNA for 14-3-3β and 14-3-3γ on the transdifferentiation potential of NIH3T3-L1 cells into osteoblasts using a retroviral gene-delivery system. The shRNAs were constructed following the guidelines from [34] and their efficiency and specificity were tested (Fig 6C). Figure 6A showed the detection of the osteoblastic early-differentiation marker, alkaline phosphatase, after treating the cells with ODM for 7 days. We observed a significantly distinctive trans-differentiation potential of NIH3T3-L1 WT, shRNA 14-3-3β or 14-3-3γ when they were induced to differentiate by ODM. While the absence of expression of 14-3-3β (Fig 6A) makes the cells more prone to differentiate to osteoblasts, the absence of expression of 14-3-3γ has the opposite effect on these cells. The quatification of this effect is shown in the figure 6B. This result is in agreement with the relatively lower levels of 14-3-3β at the beginning of the course of ASC osteoblastic differentiation (Fig 1).

**FIGURE 6.**
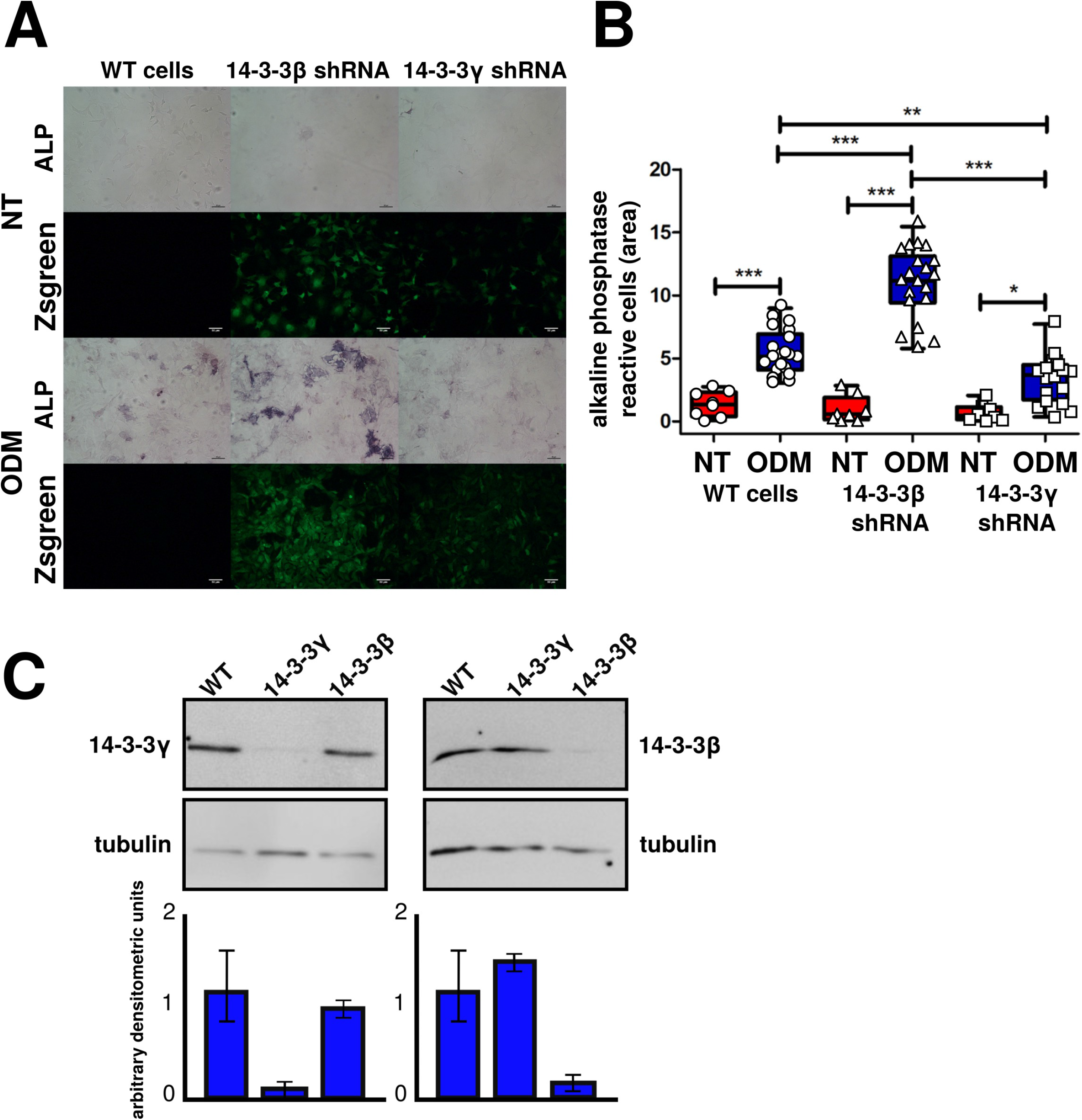
Alkaline phosphatase detection in WT and 14-3-3β or γ paralogs-silenced NIH3T3-L1 preadipocytes trans-differentiated to the osteogenic lineage. A) 14-3-3β and γ paralogs silencing. Western blot analysis of protein extracts from NIH3T3-L1 cells that where infected with shRNA-containing lentiviruses to generate 14-3-3β and γ-silenced NIH3T3-L1 stable cell lines. 40 μg were loaded per well and separated by SDS-PAGE. Blots were divided for reaction with specific antibodies against 14-3-3β and γ, and tubulin as loading control. Bars in the graphs represent mean ± SD, of values from 3 experiments quantified by densitometry and expressed relative to that of tubulin values from the same Western blot line. Asterisks indicate significant differences between samples (using the Wilcoxon rank-sum test, *p* < 0.05). **B**) Bright-field and fluorescent microscopy images (20X) of fixed NIH3T3-L1 cells (NT, non-treated) and treated with osteogenic differentiation medium (ODM) for 7 days. Cells were fixed and alkaline phosphatase activity was detected (purple staining) in NIH3T3-L1 uninfected (WT) and infected cells with lentivirus containing 14-3-3β and γ shRNAs. Green signal corresponds to the ZsGreen gene in the lentivirus particles. C) Quantification of alkaline phosphatase reactive cells (purple area) in NT and ODM, uninfected (WT) and infected (14-3-3β and γ shRNAs) NIH3T3-L1 cells. Bars represent mean values (± standard deviation) from 10 stained images. The asterisks indicate significant differences between samples (Wilcoxon rank-sum test, *p* < 0.01). Data were similar in two other experiments.

Our study demonstrated an important regulatory implication of 14-3-3β (negative) and 14-3-3γ (positive) in the transdifferentiation of preadipocytes into differentiated bone-forming osteoblasts, suppressing adipogenesis of NIH3T3-L1 preadipocytes and stimulating osteoblastic differentiation.

## Discussion

Numerous examples show that the post-translational modification of N-term of histone by acetylation is highly relevant in many cells, including ASC [35]. Defects on this modification frequently result in severe abnormalities of development, differentiation and physiology, mostly due to an aberrant gene expression [36]. Most mammalian proteins are modified by more than one PTMs, which usually influence each other, phenomenon often referred to as PTM crosstalk. Transcription factors and other transcriptional regulators are involved in the most extensively studied PTM crosstalk with acetylated non-histone proteins [36]. Although there are several identified interactions between non-histone protein acetylation and protein phosphorylation signaling [12], the case of the 14-3-3 proteins [14] is different. The 14-3-3 protein family is considered as a master phosphorylation reader [16]. They can bind to pSer/Thr in their target proteins, and recently were identified to have several Lys residues modified by acetylation [14]. Among them, the Lys49 (or Lys51) is essential for the binding to their target proteins because it forms one of the three hydrogen bond needed for the interaction to occur [37]. Andersen and coworkers identified the cytoplasmic deacetylase HDAC6 as the responsible to eliminate the acetyl group from the acetyl-N-ε-Lys49/51 [15]. The acetylation of 14-3-3ζ on this residue blocks the interactions with Bad and AS160, two well described interacting partners of 14-3-3 proteins [15], and to a phosphorylated RAF-1 peptide [14]. Once the binding to 14-3-3 is impaired, the Ser112 of BAD and the Thr642 of AS160 are much more accessible for their dephosphorylation [15], demonstrating the control of 14-3-3 activity by the reversible cycle of acetylation/deacetylation. This kind of regulation was theoretically proposed earlier by Beltrao and coworkers, who named it as “regulation of regulators” [16]. Our results show that the acetyltransferase HBO1 (KAT7) modifies 14-3-3 proteins (probably all paralogs) in the Lys49/51. The HBO1 subcellular localization is mostly nuclear [36], althought our results of confocal microscopy showed some level of immunoreactivity in the cytoplasm (Fig 2 and others). Zou and co-workers also detected functional HBO1 in the cytoplasm [28]. Among acetyltransferases, only the highly specific α-tubulin *N*-acetyltransferase 1 is detected in the cytoplasm [38]. The cytoplasmatic deacetylase activity is also highly restricted, being only the class IIb (the class where the previously mentioned 14-3-3 deacetylase HDAC6 belongs) present in this compartment, whereas the class I, IIa and IV are exclusively nuclear [36]. These subcellular localizations of the HBO1 acetyltransferase and the HDAC6 deacetylase are in agreement with our results that show high levels of immunoreactivity against 14-3-3 acK49/51 in the nucleus, where this residue could be only modified by the HBO1, and lower levels in the cytoplasm where both activities are present. Looking for evidence of binding between 14-3-3 proteins and members of acetyltransferases families, only KAT7 (HBO1) and KAT8 have the potential to interact with 14-3-3s proteins (https://ania-1433.lifesci.dundee.ac.uk/prediction/webserver/index.py) [39].

Inhibition of the activity of HDACs with specific or pan HDAC inhibitors, enhances osteogenic differentiation in detriment of adipogenic differentiation in many cellular models [6,30,40]. These effects could be a consequence of the change of the acetylation state in the N-term of histone H3 and/or H4. Our results show that 14-3-3β acetylation is also involved in osteogenic differentiation. The acetylation of 14-3-3 acK49/51 produces its inhibition and we observed that during the osteogenic differentiation the inhibition of 14-3-3 (specifically 14-3-3β) increased. It has been demonstrated that the glycerol-phosphate inhibits the 14-3-3 proteins [41], and the use of the later in the osteogenic differentiation drug cocktail reinforces the idea that inhibition of 14-3-3 proteins is necessary for this differentiation to occur *in vitro.* 14-3-3β was described as a negative regulator of osteogenesis [17] but the specific mechanism involved remains unclear. The integration of our results could explain that the site-specific inhibition of 14-3-3β by HBO1 acetylation on Lys49/51 reduces its activity. This could alter the subcellular localization of its binding partners like the transcriptional coactivator with PDZ-binding motif, TAZ, a modulator of mesenchymal stem cells osteogenic differentiation [19]. Transdifferentiation of NIH3T3-L1 cells in which the specific expression of 14-3-3β or 14-3-3γ were reduced showed differences in their potential to differentiate to osteoblasts (Fig 6). To our knowledge, this is the first time in which the acetylation of one paralog of the 14-3-3 family is linked to a specific biological role.

## Materials and Methods

### Cell Culture

Healthy patients who underwent a dermolipectomy voluntarily donated abdominal subcutaneous adipose tissue after signing an informed consent in accordance with the Declaration of Helsinki (*Universidad Nacional de Cuyo* Bioethics Committee 14594/2014). The material was processed to obtain the ASCs from the stromal vascular fraction as described in [21]. Cells were characterized by flow cytometry, cryopreserved and used between passages 4 to 9.

The ASCs as well as the NIH3T3-L1 preadipocyte cell line (ATCC, United States) were maintained in high glucose Dulbecco’s Modified Eagle Medium (DMEM, *Thermo Fisher)* supplemented with 10% Fetal Bovine Serum (FBS-*Internegocios S.A.,* Buenos Aires, Argentina), L-Glutamine 1%, and antibiotics penicillin 10 U/mL and streptomycin 100 μg/mL *(Thermo Fisher)* at 37°C in a 5% CO_2_ atmosphere.

### SDS-PAGE and Western blotting

Cells were lyzed using RIPA buffer and proteins were quantified using the BCA method (Protein BCA Assay Kit, *Thermo Fisher).* The protein extracts were separeted on 12% SDS-PAGE gels (50 μg of total proteins were loaded per well) and then electro-transferred to polyvinylidene fluoride (PVDF) membranes. The membranes were blocked in 5% skimmed milk in PBS for 30 min at RT, and then incubated with primary antibodies for 1 h at RT. Membranes were washed with PBST 0,01%, then probed with HRP-conjugated goat anti-rabbit IgG or horse anti-mouse IgG secondary antibodies diluted at 1:5000 in 5% skimmed milk in PBST 0,01% for 1 h at RT. Finally, the membranes were washed three times with PBST 0,01% and developed using the ECL kit. Band intensities were quantified by the densitometric analysis software Image Studio™ Lite 5.2 (LI-COR Biosciences). Relative expression levels of target proteins were normalized to β-tubulin used as loading control. All the experiments were repeated three independent times.

### Antibodies

Antibodies used for Western blotting, immunostaining and immunoprecipitation included anti-14-3-3β; anti-14-3-3ε; anti-14-3-3η; anti-14-3-3γ all this from [32]; anti-14-3-3τ (sc-732, Santa Cruz Biotechnology), anti-14-3-3ζ (1019, Santa Cruz Biochnology); anti-14-3-3 acK49/51 (ST97695, St. John’s Laboratory); anti-RUNX2 (390351, Santa Cruz Biotechnology); anti-HBO1 (388346, Santa Cruz Biotechnology); anti-β-tubulin (T4026, Sigma-Aldrich); Goat anti-rabbit IgG-HRP (W401B, Promega); Horse anti-mouse IgG-HRP (PI2000, Vector Lab); Donkey anti-mouse IgG Alexa 594 (A-21203, Thermo Fisher); Goat anti-Rabbit IgG Alexa Fluor 488 (A-11008, Thermo Fisher); preimmune Rabbit Serum.

### Osteogenic differentiation of ASCs

The ASCs were seeded at a density of 2,5.10^4^ cells/cm^2^ and after 72 h were induced to differentiate to osteoblasts by adding ODM containing DMEM with low glucose (Thermo Fisher) supplemented with 10% FBS (Internegocios); L-Glutamine 1%; β-glycerophosphate (β-GP – Sigma-Aldrich) 10 mM; 2-phospho-L-ascorbic acid (AA2P – *Sigma-Aldrich)* 50 μg/μL; dexamethasone (Sigma-Aldrich) 0.1 μM and antibiotics penicillin 10 U/mL and streptomycin 100 μg/mL (Thermo Fisher) at 37°C in a 5% CO2 atmosphere during the specified periods with medium changes every two or three days. Negative controls were maintained in DMEM with low glucose supplemented with 10% FBS.

### Lentiviral shRNA knockdown of 14-3-3β and 14-3-3γ in NIH3T3-L1 cell line

Plasmids encoding lentiviruses expressing shRNAs were designed following guide lines from [34]. Plasmids were first purified with a midiprep kit (Roche) and were transfected into HEK293T cells with a five-plasmid system to produce lentivirus with a very high titer of ~10^7^ CFU/ml. 14-3-3β shRNA (tgctgttgacagtgagcgcgctctctgttgcctacaagaatagtgaagccacagatgtattcttgtaggcaacagagagcatgcctactgcctcgga), and 14-3-3γ shRNA (gctactactgcagtctttatgctgttgacagtgagcgctgctactactgcagtctttattagtgaagccacagatgtaataaagactgcagtagtagcattgcctactgcctcgga) were produced. As these lentiviruses simultaneously express ZsGreen by an IRES system, the levels of infection were checked by flow cytometry analyses. We routinely obtained more than 90% of positive cells.

### Transdifferentiation of NIH3T3-L1 cell line

The NIH3T3-L1 cell lines (WT,14-3-3β and 14-3-3γ-silenced) were seeded onto 24-well plates at a cell density of 12,5.10^3^ cells/cm^2^, and after 24 h were induced to transdifferentiate by adding ODM as specified above for ASC osteoblastic differentiation, but without dexamethasone [42]. After that, medium was changed every two or three days by ODM with dexametasone 10 nM. Negative controls were maintained the first three days in DMEM with low glucose supplemented with 5% FBS, and the following days with DMEM supplemented with 2,5% FBS.

### Osteogenic specific linage stainings

ASCs differentiated to osteoblasts and their corresponding negative controls were stained with Alizarin Red Solution (Biopack) at different days after induction of differentiation. Briefly, the cells adhered to circular 12 mm diameter coverslips were washed twice with PBS and then fixed with 4% paraformaldehyde (PAF) for 20 min. The fixed cells were washed twice with PBS and incubated at room temperature, in darkness and constant agitation for 20 min with ARS 40 mM (pH 4.1-4.3). The cells were thoroughly washed with Milli-Q water to remove dye excess, and the coverslips were mounted on slides with Mowiol 4-88 for analysis by optical microscopy.

For alkaline phosphatase (ALP) staining, cells grown on circular 12 mm diameter coverslips were washed with PBS twice and then fixed with 4% PAF for 5 min. Subsequently, the cells were permeabilized twice with PBS 0.05% Tween-20 (PBST) for 2.5 min. Then, the permeabilization buffer was aspirated and the coverslips were incubated for 1 h with 500 μL *per* well of 100 mM Tris-HCl (pH 8.30), 5 mM MgCl_2_ reaction buffer containing 0.015% w/v 5-bromo-4-chloro-3-indolyl phosphate-BCIP-(*Calbiochem)* substrate and 0.03% w/v nitro blue tetrazolium-NBT-*(BDH Chemicals Ltd.)* at room temperature, in darkness and with constant agitation. After exhaust washing, the coverslips were mounted with Mowiol 4-88 for further analysis by optical microscopy.

ALP kinetics was measured using a soluble and colorless synthetic substrate, p-nitrophenylphosphate-pNPP-(Calbiochem, San Diego, CA, USA), whose hydrolysis generates a yellow product, *p*-nitrophenol. Briefly, the cells were fixed with 4% PFA for 5 min, permeabilized with PBST for 5 min, and the monolayers were washed with 50 mM bicarbonate buffer (pH 9.60) and 1 mM MgCl_2_. Subsequently, 1 mL of bicarbonate buffer containing 10 mM pNPP was added to each well, and the plate was incubated at 30°C with gentle agitation for 90 min, taking a 100 μL sample per well every 10 min for one hour, and then a final sample after 90 min. Each sample was immediately transferred to a 96-well plate previously loaded with 50 μL 3N NaOH to stop the reaction. At the end of the sampling, absorbance at 405 nm was measured in a MultiSkan FC microplate reader (Thermo Fisher Scientific).

### Treatment with sodium butyrate and lactacystin

The ASCs were treated with sodium butyrate (NaBu – Sigma-Aldrich) solution in H_2_O Milli-Q at a final concentration of 0.5 mM for three days before osteogenic differentiation induction.

Lactacystin (Lac-L6785, Sigma-Aldrich) solution in H_2_O Milli-Q was added to the culture media at a concentration of 10 μM for 24 h before performing the ASCs indirect immunofluorescent assay at different days (6, 13 and 20) after induction to osteoblasts differentiation.

### Immunoprecipitation of 14-3-3β and 14-3-3γ

The ASCs on day 21 (d21) after induction of differentiation were lysed in lysis buffer (50 mM Tris-HCl pH 6.8; Nonidet P-40 1% v/v; SDS 0.05% w/v; 2 mM EDTA, containing deactylase and protease inhibitors). Cell lysates (250 μg total proteins) were subjected to immunoprecipitation with anti-14-3-3β [32], anti-14-3-3γ [32] and preimmune rabbit serum (5 μL of each sera), and proteins precipitated using protein-A Agarose (Sigma-Aldrich) prior to analyses by Western blotting.

### Indirect immunofluorescence

The ASCs grown on 12 mm diameter coverslips on days 7, 14 and 21 (d7, d14 and d21, respectively) after differentiation induction were washed twice with PBS supplemented with 1 mM CaCl_2_ and MgCl_2_, fixed with 4% PAF for 20 min and permeabilized with 0.3% Triton X-100 in PBS, and blocked in 10% BSA-PBS prior to addition of antibodies. Primary antibodies used were mouse anti-HBO1 (sc-388346, *Santa Cruz Biotechnology Inc.,* 1:100); rabbit anti-14-3-3 acK49/51 (SJ97695, St. John’s Laboratory Ltd., 1:200), and secondary antibodies were Donkey anti-mouse IgG Alexa 594 (A-21203, Thermo Fisher, 1:250); Goat anti-Rabbit IgG Alexa Fluor 488 (A-11008, Thermo Fisher, 1:200) in 2% BSA-PBS. Cell nuclei were visualized by DAPI (Vector) staining. Images were captured using a confocal microscope Olympus FluoView 1000. For quantification of 14-3-3 acK49/51 localization, a minimum of five images on each condition and nuclear/cytoplasmic localization of 14-3-3 acK49/51 was processed using the ImageJ software or in house Python scripts.

### Statistical and automatic confocal image analysis

The statistical test performed at each experiment, as well n and *p* value obtained is indicated in the figure legends. The Pearson, Manders and Li’s correlation coefficients for the colocalization were determined by using the ImageJ software [43] plugings JACoP [44]. The GraphPad Prism 5 software or Rstudio were used for the analysis.

## Acknowledgements

YRFL and ADG are fellows from CONICET; MU and DMB are members of the Investigator Career of the National Research Council (CONICET, Argentina). We acknowledge the ANPCyT (PICT’14 1659, PICT’17 1984), MINCyT DAAD HA16/02 and Fundación Roemmers for funding the project. The funding institutions had no role in study design, analysis and decision to publish. The lentiviral system was kindly provided by Dr. Gustavo Mostoslavsky (Boston University), as well as reagents and enzymes used for their cloning during an ASBMB fellowship granted to ADG. The plasmid pCMV-HBO1-3xTag-6 was kindly provide by Dr. Yeun Kyu Jang (Yonsei University, South Korea). We thank Jorge Ibañez (IHEM) for the technical support in confocal microscopy.

## Author contributions

YRFL and ADG perform the experiments. YRFL, MU, and DMB analyzed, quantified and interpreted the data. YRFL, MU, and DMB contributed to the conception and design of the study. YRFL, MU, and DMB wrote the manuscript and provided critical revisions. All authors approved the final manuscript.

---

**FIGURE S1.**
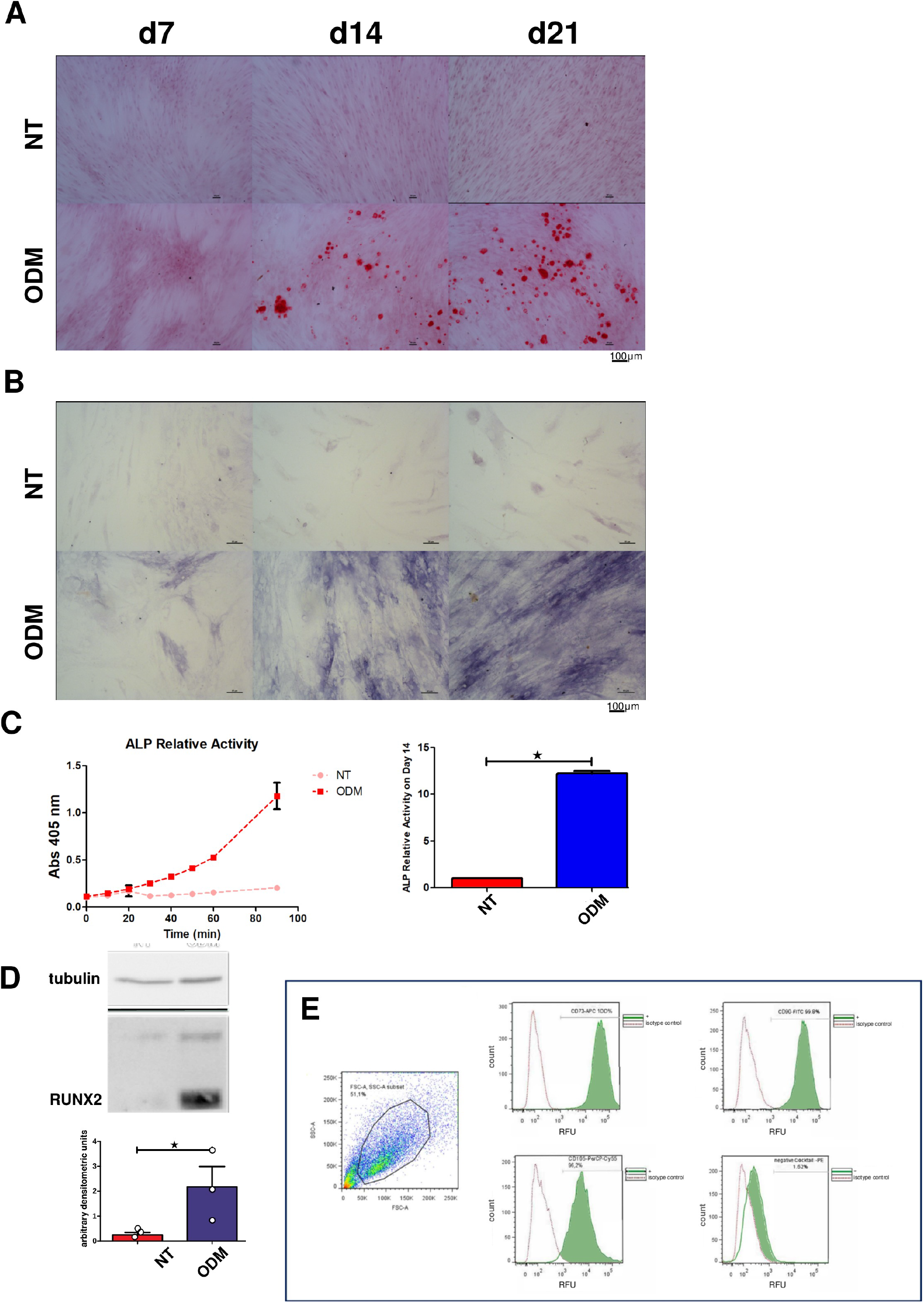
Osteogenic differentiation of ASC. A, B) Bright-field microscopy images of fixed ASC (NT, nontreated) and treated with osteogenic differentiation medium (ODM) as described in the methods section. A) Alizarin red staining images (magnification 10X) at 7, 14 and 21 days after induction. B) Same as in B except that cells were stained for alkaline phosphatase detection (magnification 20X). C) Soluble alkaline phosphatase activity quantification through absorbance of *p*-nitrophenol at 405 nm in non-treated (NT) and ODM-treated ASC for 14 days. D) Western blot analysis of RUNX2 protein levels in non-treated (NT) and ODM-treated ASC for 21 days. Blots were prepared as in figure 1 and incubated with antibodies against RUNX2, and tubulin as loading control. Graphs were constructed as in figure 1 except that RUNX2 densitometry values were expressed relative to that of tubulin. E) Multicolor phenotyping of ASCs by flow citometry. Histograms represent the number of events (count) vs. the relative fluorescence units (RFU) for each indicated fluorophore conjugated-antibody or cocktail. Three positive (CD105, CD90 and CD73) and five negative (CD45, CD34, CD11b, CD19 and HLA-DR) markers. A to E) Data were similar in two other experiments.

**FIGURE S2.**
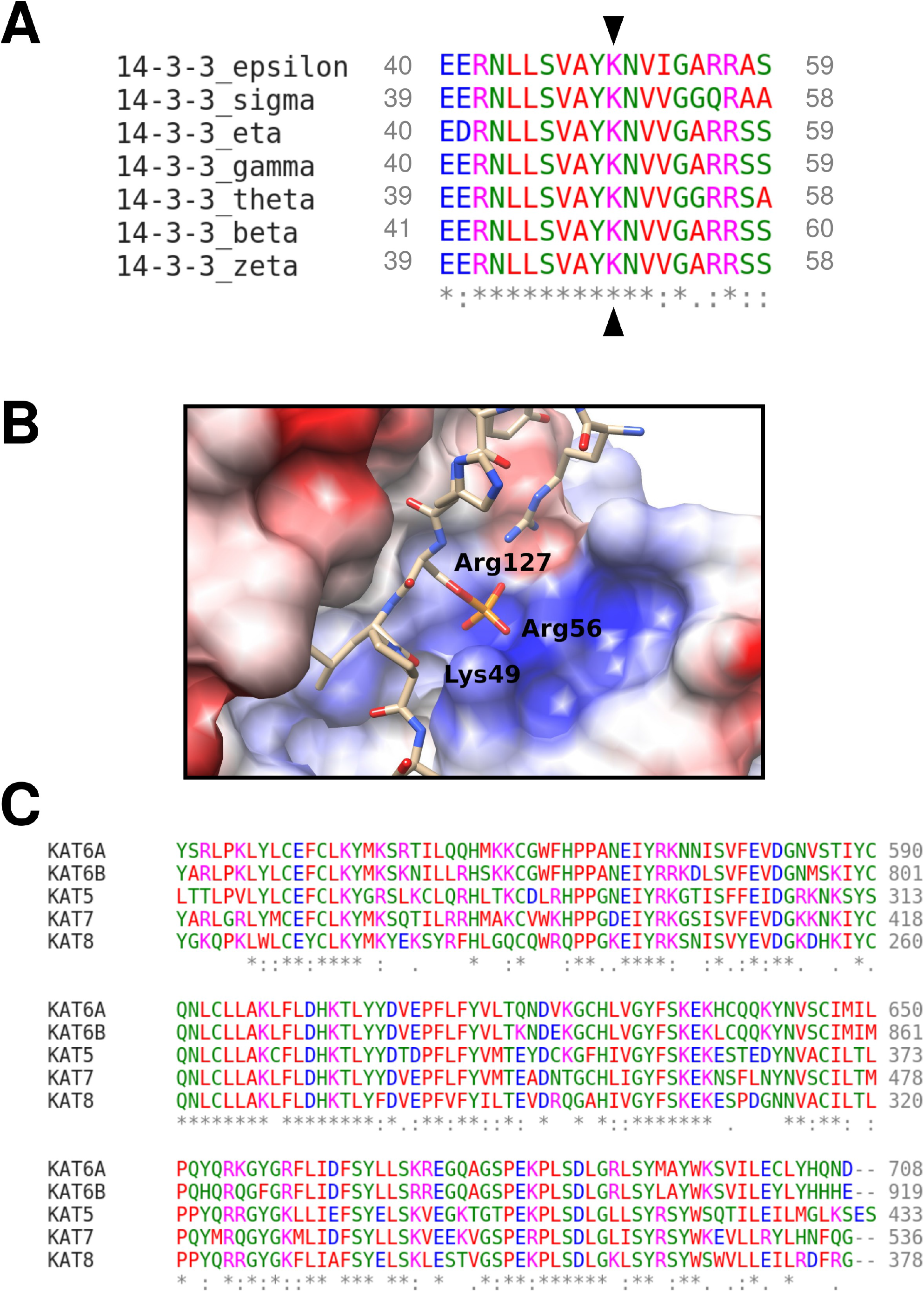
Lys49/51 (arrows) in the 14-3-3 protein family and Myst family alignment. A) Protein sequences alignment comparing the K49/51 and neighboring residues in the seven 14-3-3 mammalian paralogs. B) X-ray crystallographic structure of 14-3-3 binding pocket (1QJA). 14-3-3ζ is bound to a phosphoserine peptides. The critical residues Lys 49, Arg 56 and 127 in 14-3-3 protein are shown. Blue, positive charges; red, negative charges. The phosphoserine from the peptide is close to the three essential residues in 14-3-3 forming the hydrogen bonds necessary for the binding. C) Protein sequences alignment of histone deacetlylases belonging to the MYST family. KAT7 is a synonym of HBO1.

**FIGURE S3.**
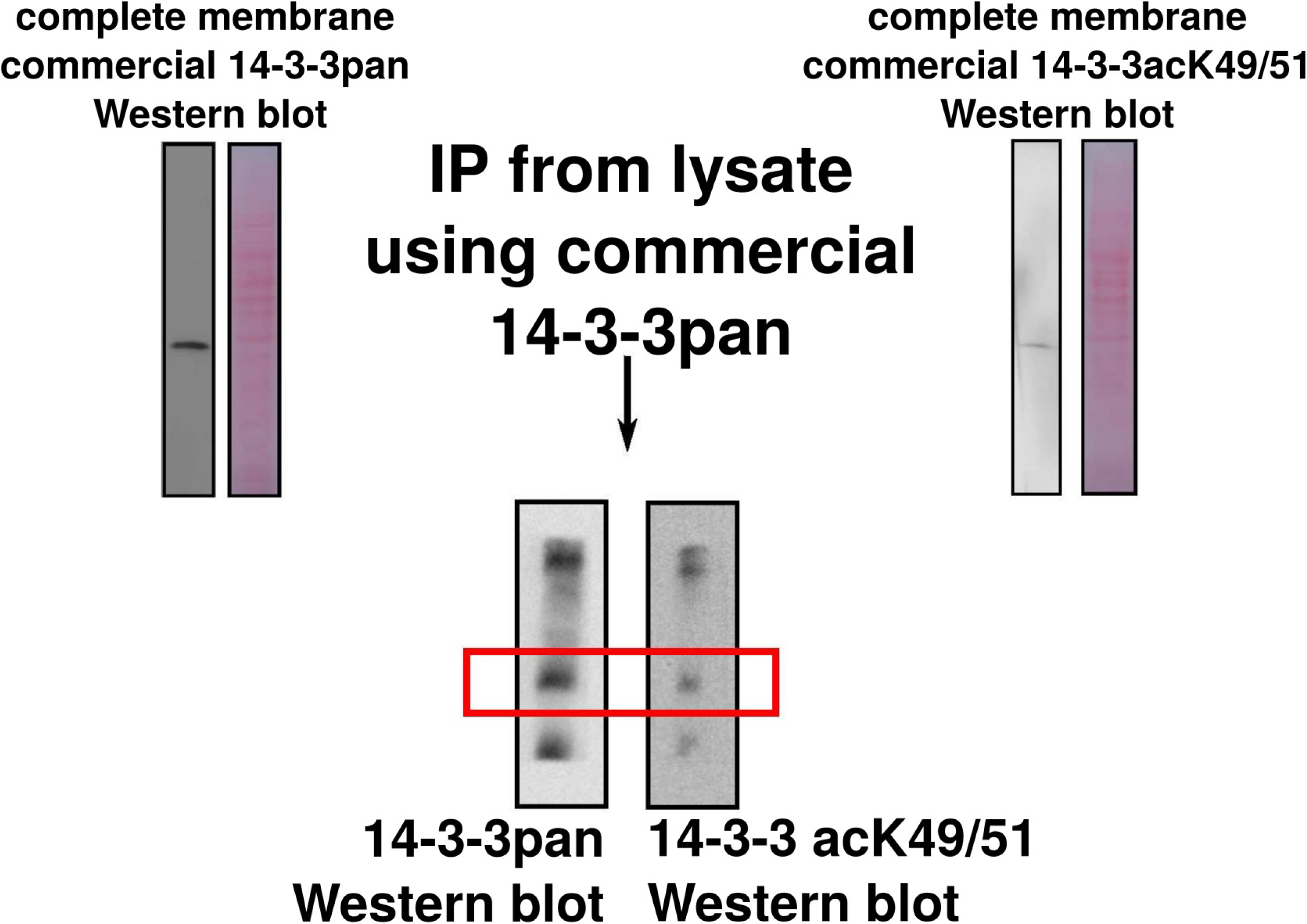
Characterization of 14-3-3 acK49/51 antibody. Antibody ST97695 from St. John’s Laboratory against 14-3-3acK49/51 were characterized by immunoprecipitation of 14-3-3 protein from ASC lysates following by SDS-PAGE and Western blot. Also the company provide us with the acetylated peptide used in ELISA experiments to characterize the antibody EERNLLSVAY(ac)KNVVG.

**FIGURE S4.**
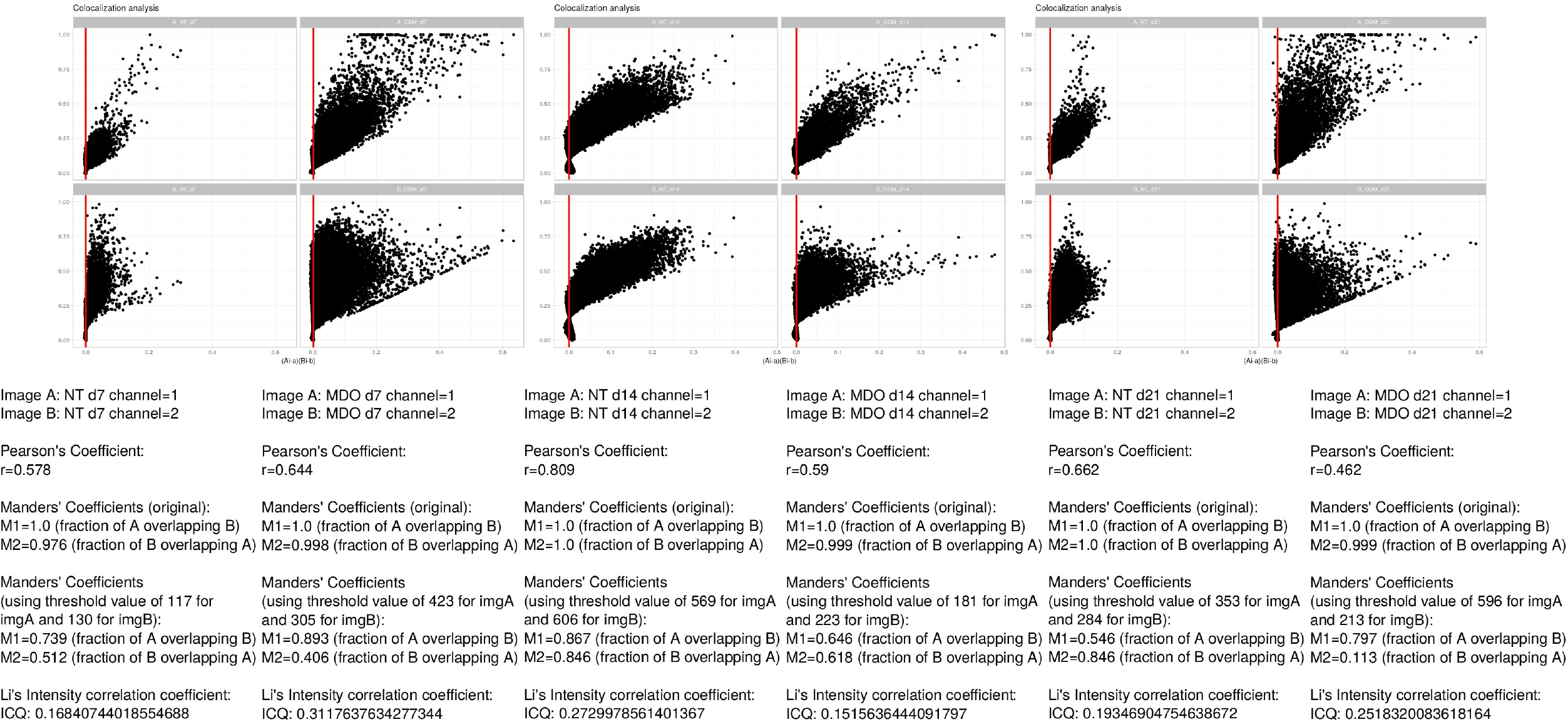
Analysis of 14-3-3 acK49/51 and HBO1 co-localization. Inset images from figure 3 (or similar images from other experiments) were analyzed by Jacop plugin [44] of ImageJ. This package do quantitative colocalization of confocal images by using standard methods (Pearson, Manders, corrected Manders and Li). All parameters were maintained as default.

**FIGURE S5.**
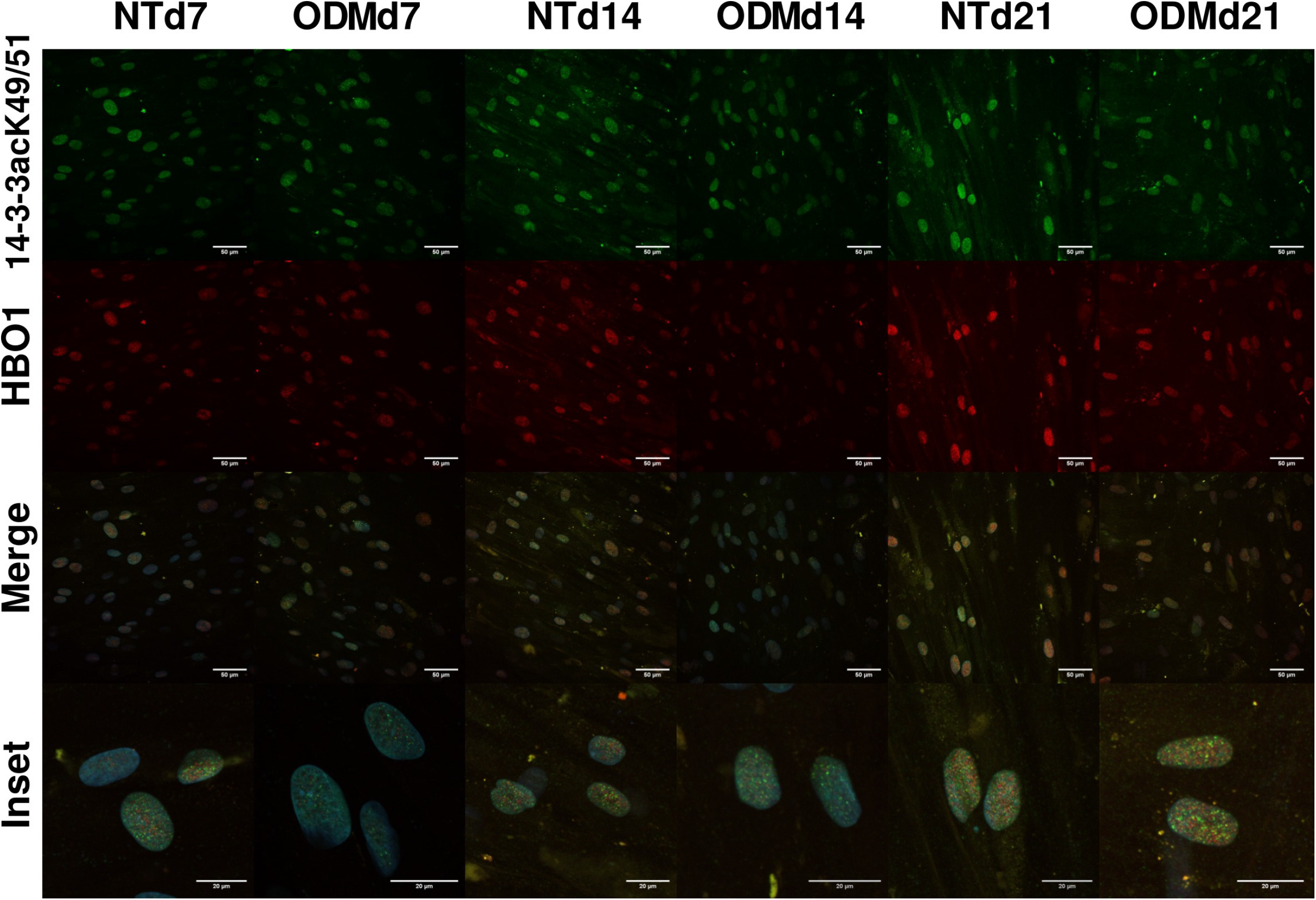
14-3-3 acK49/51 and HBO1 in Lactacystin treated cells. Same as Fig. 3 except that ASC were treated with 10 μM Lactacystin for 24 h before performing the ASCs indirect immunofluorescent assay at different days (6, 13 and 20) after induction to osteoblasts differentiation. Magnification 40X. Data were similar in two other experiments.

